# Stabilin levels are related to atherosclerotic plaque burden in type 2 diabetes mellitus individuals

**DOI:** 10.1101/2025.07.10.664255

**Authors:** Eirini Giannousi, Chara Georgiadou, Nikolaos I. Vlachogiannis, Eva Kassi, Ioanna-Katerina Aggeli, Petros P. Sfikakis, Nikolaos Tentolouris, Athanase D. Protogerou, Georgios Kararigas, Antonios Chatzigeorgiou

## Abstract

Type 2 diabetes mellitus (T2DM) is a global burgeoning health problem that increases the risk of atherosclerotic cardiovascular disease (ASCVD). Infiltration and oxidative modification of low-density lipoprotein (LDL) cholesterol in the arterial wall and chronic inflammation comprise central pathogenetic mechanisms in ASCVD. Scavenger receptors, particularly Stabilin-1 (Stab1) and Stabilin-2 (Stab2), are pivotal in the clearance of oxidized LDL (oxLDL) cholesterol and pro-atherogenic ligands from circulation. However, their role in atherosclerosis development in the spectrum of T2DM remains poorly characterized. We assessed circulating levels of Stab1, Stab2, and their ligands (TGFbI, Periostin and Reelin) in a cohort of 33 T2DM and 21 non-diabetic individuals, stratified by their atherosclerotic plaque burden as assessed by high-resolution vascular ultrasound. Associations between stabilins, their ligands and conventional cardiovascular risk factors were evaluated. Stab1 levels were significantly elevated in individuals with higher atherosclerotic plaque burden (p<0.05), while Reelin levels were marginally elevated, both in the total study cohort and among T2DM patients. Stab1 levels positively correlated with body mass index and inversely correlated with total cholesterol, LDL, and high-density lipoprotein (HDL) cholesterol levels. Our findings indicate that Stab1 may serve as a marker of dysregulated lipid metabolism and increased atherosclerotic plaque burden in individuals with T2DM. Larger prospective studies are warranted to establish the prognostic and potentially therapeutic value of Stab1 and to clarify its mechanistic role in diabetic atherosclerosis.

## Introduction

Type 2 diabetes mellitus (T2DM) represents a growing global health pandemic, with projections indicating a substantial rise in prevalence over coming decades^1^. Individuals with T2DM face significantly elevated risks of cardiovascular disease (CVD), which accounts for 40-60% of mortality in this population^2–4^. Atherosclerotic CVD (ASCVD), a common manifestation of CVD, is a major cause of mortality in diabetic patients^5^. ASCVD arises from the extensive recruitment and deposition of serum lipoproteins in the arterial wall, driven by complex interactions between metabolic dysregulation, endothelial dysfunction and chronic inflammation in T2DM^6–8^. Numerous factors contribute to the development and progression of ASCVD, including hemostatic factors, lipoprotein metabolism, insulin resistance, chronic inflammation, oxidative stress, and endothelial dysfunction^5,9–11^. Despite advances in lipid-lowering medications, atherosclerosis remains responsible for the majority of cardiovascular deaths, emphasizing an urgent clinical need to uncover novel therapeutic targets for ASCVD^12,13^.

In this context, scavenger receptors (SR) constitute a category of potential target proteins. SR mediate the uptake of oxidized low-density lipoprotein (oxLDL) cholesterol by macrophages and endothelial cells in the sub-endothelial region of the artery^14,15^. Notably, Stabilin-1 (Stab1) and Stabilin-2 (Stab2), are SR expressed mainly on liver sinusoidal endothelial cells (LSECs) and have the ability to bind to several ligands, including oxLDL, acetylated LDL, advanced glycation end products, and collagen propeptides^16–18^. Stabilins affect atherogenesis through the clearance of pro-atherogenic ligands, such as Reelin, Periostin, TGFbI, and suppression of monocyte/macrophage activation, thus attenuating inflammatory cascades^12,19,20^. Of note, a recent preclinical study showed that in both spontaneous and western diet-induced models of atherosclerosis, aortic plaque growth was markedly decreased by both genetic deletion and antibody targeting of Stab1 and Stab2^12^. In the present study, we aimed at exploring the potential value of Stabilins 1 and 2 as non-invasive markers of atherosclerotic plaque burden in individuals with or without T2DM.

## Methods

### Patients, Biochemical Parameters and Atherosclerosis Assessment

In this study, 54 individuals were included; 33 individuals with T2DM and 21 non-diabetic individuals of similar age and sex (Table 1). All individuals enrolled in the study had no history of CVD, chronic renal disease, or any chronic inflammatory disease at the time of inclusion in the study. The study was approved by the Laiko Hospital Ethics Committee and informed consent was obtained from all participants in accordance with the Declaration of Helsinki. All study participants underwent blood sampling after overnight fasting. Detailed medical history was collected for every individual, including anthropometric assessment, history of CVD risk factors, and detailed biochemistry profile of liver and renal function markers, markers of systemic inflammation and cholesterol levels. Blood pressure (BP) was measured twice in both arms while participants were in the supine position, with measurements taken by a trained physician. The estimated glomerular filtration rate (eGFR) was calculated with the CKD-EPI equation^21^. Further methodological details including the evaluation for the presence of atheromatic plaques can be found in previous works^22,23^. The 54 patients included in the study were categorized based on their atherosclerotic burden (0–1 number of atherosclerotic plaques; ≥2 number of atherosclerotic plaques).

**Table 1.** Demographics and clinical characteristics of the study cohort.

|  |  |  |  |  |
| --- | --- | --- | --- | --- |
|  | **Whole cohort**<br>**(n=54)** | **Non-diabetic**<br>**(n=21)** | **T2DM**<br>**(n=33)** | **P-value\*** |
| Age (years) | 65.3±7.7 | 65.8±7.3 | 65.0±8.1 | 0.728 |
| Male sex | 24(44.4) | 9 (42.9) | 15 (45.5) | 1.000 |
| **BMI (kg/m<sup>2</sup>)** | **31.2±6.5** | **28.0±4.4** | **33.2±6.9** | **0.002** |
| Smoking (active) | 8 (14.8) | 4 (19.0) | 4 (12.1) | 0.697 |
| Arterial hypertension | 45 (83.3) | 18 (85.7) | 27 (81.8) | 1.000 |
| Hyperlipidemia | 38 (70.4) | 14 (66.7) | 24 (72.7) | 0.762 |
| **SBP (mmHg)** | **137±21** | **130±17** | **141±22** | **0.057** |
| DBP (mmHg) | 78±10 | 78±7 | 79±12 | 0.936** |
| **Pulse pressure (mmHg)** | **58±18** | **52±13** | **62±20** | **0.047** |
| **Total cholesterol (mg/dL)** | **193±39** | **210±39** | **183±36** | **0.012** |
| **LDL (mg/dL)** | **115±30** | **128±33** | **107±25** | **0.013** |
| **HDL (mg/dL)** | **54±15** | **61±17** | **49±12** | **0.009** |
| Triglycerides (mg/dL) | 122±68 | 108±59 | 125±59 | 0.086** |
| HbA1c (%) |  | n/a | 7.2±1.0 |  |
| eGFR (mL/min/1.73m<sup>2</sup>) | 83±20 (n=50) | 84±18 (n=18) | 83±21 | 0.936** |
| 0-1 / ≥2 atherosclerotic plaques | 28 / 26 | 10 / 11 | 18 / 15 | 0.781 |
| **Therapy** |  |  |  |  |
| Insulin | 15 (27.8) | 0 | 15 (45.5) | <0.001 |
| Anti-hypertensive drugs | 44 (81.5) | 17 (81.0) | 27 (81.8) | 1.000 |
| Lipid modifying drugs | 32 (59.3) | 11 (52.4) | 21 (63.6) | 0.571 |
| Continuous variables are presented as mean ± standard deviation. Categorical values are presented as absolute count (percentage). |  |  |  |  |
| *P-value corresponds to the comparison between individuals with T2DM and non-diabetic. Continuous variables were compared using independent samples t-test unless otherwise specified. Categorical variables were compared using Fisher's exact test. |  |  |  |  |
| **P-value derived by Mann Whitney test |  |  |  |  |

### Elisa Assays

Plasma concentrations of circulating TGFbI (E-EL-H1657, Elabscience), POSTN/OSF-2 (Periostin, E-EL-H6160, Elabscience), Reelin (EH2121, FineTest), Stabilin-1 (EH1574, Finetest) and Stabilin-2 (HUEB0677, AssayGenie) were measured according to manufacturers’ instructions.

### Statistical Analysis

For the statistical analysis of the data, normality of continuous variables was assessed by utilizing the Shapiro-Wilk test. The presence of outliers was examined with Rout test (Q = 1%). Comparison of continuous variables between two groups was performed by independent samples t-test with or without Welch’s correction or Mann-Whitney U test when data were non-normally distributed. Correlation between continuous variables was examined by Spearman’s rank test. The statistical analysis was conducted utilizing Graph Pad Prism 10 and IBM SPSS Statistics v.30. Statistical significance was set at p < 0.05.

## Results

### Key clinical characteristics of non-diabetic and T2DM individuals

The present study cohort comprised 54 individuals, including 21 non-diabetic and 33 T2DM individuals (Table 1). The age and the proportion of male participants were similar in both groups (non-diabetic vs. T2DM: Age: 65.8 ± 7.3 vs. 65.0 ± 8.1 years, p = 0.728; Male sex: 42.9% vs 45.5%. p = 1.000). Body mass index (BMI) was significantly higher in the T2DM compared to the non-diabetic group (non-diabetic vs. T2DM: 28.0 ± 4.4 vs. 33.2 ± 6.9 kg/m^2^, p = 0.002). Systolic blood pressure (SBP) and pulse pressure tended to be higher in T2DM patients (non-diabetic vs. T2DM: SBP: 130 ± 17 vs. 141 ± 22 mmHg, p = 0.057; pulse pressure: 52 ± 13 vs. 62 ± 20 mmHg, p = 0.047). T2DM patients exhibited significantly lower total cholesterol (210 ± 39 vs 183 ± 36 mg/dL, p = 0.012), LDL cholesterol (128 ± 33 vs. 107 ± 25 mg/dL, p = 0.013), and high-density lipoprotein (HDL) cholesterol (61 ± 17 vs. 49 ± 12 mg/dL, p = 0.009) compared to the non-diabetic groups. The distribution of atherosclerotic plaque burden (0–1 vs. ≥2 atherosclerotic plaques) did not differ between non-diabetic and T2DM individuals.

### Increased circulating levels of Stabilin-1 in patients with high atherosclerotic burden

Circulating levels of TGFbI, Periostin, Reelin, Stab1 and Stab2 were measured in the pooled cohort of non-diabetic and T2DM individuals, stratified by the number of atherosclerotic plaques. The circulating levels of the ligands TGFbI, Periostin and Reelin were comparable between the two groups (low vs. high atherosclerotic burden: TGFbI: 577.6 ± 57.61 vs. 684 ± 67.76 ng/ml, p = 0.3; Periostin: 108.6 ± 9.985 vs. 128.3 ± 14.42 ng/ml, p = 0.507; Reelin: 1637 ± 201.5 vs. 2318 ± 324 pg/ml, p = 0.109; Fig. 1A-C). Importantly, Reelin was below the detection limit of the assay in 13/54 individuals. On the other hand, Stab1 levels were significantly elevated in patients with a higher number of atherosclerotic plaques (low vs. high atherosclerotic burden: Stab1: 18.33 ±1.476 vs. 27.17 ± 2.652 ng/ml, p = 0.016; Fig. 1D), while Stab2 levels were comparable between the two groups (low vs. high atherosclerotic burden: Stab2: 3963 ± 335.5 vs. 4259 ± 404.4 pg/ml, p = 0.75; Fig. 1E).

**Figure 1.**
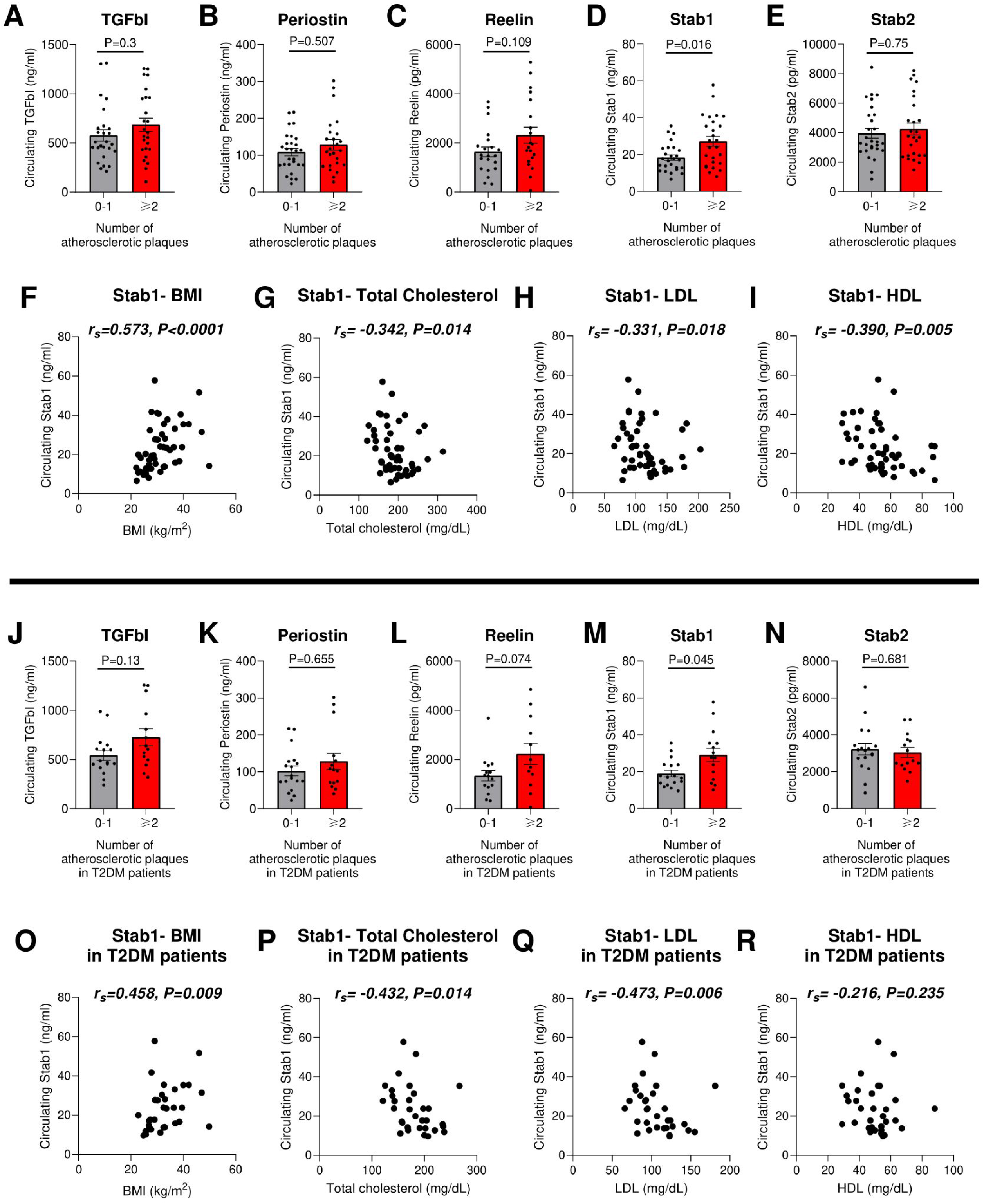
Levels of receptors Stabilin 1 & 2, their ligands TGFbI, Periostin and Reelin and correlation of Stab1 with clinical and laboratory features of atherosclerosis. **A-E)** Bar-charts showing the circulating levels of TGFbI (**A**), Periostin (**B**), Reelin (**C**), Stab1 (**D**) and Stab2 (**E**) in the plasma of nondiabetic individuals and patients with type T2DM categorized based on the number of their atherosclerotic plaques. **F-I)** Scatter-plot showing the correlation of Stab1 with BMI (**F**), total cholesterol (**G**), LDL (**H**) and HDL (**I**). **J-N)** Bar-charts showing the circulating levels of TGFbI (**J**), Periostin (**K**), Reelin (**L**), Stab1 (**M**) and Stab2 (**N**) in the plasma of T2DM patients stratified based on the number of their atherosclerotic plaques. **O-R)** Scatter-plots showing the correlation of Stab1 with BMI (**O**), total cholesterol (**P**), LDL (**Q**) and HDL (**R**). Data presented as mean ± SEM. p-Values were derived by Mann–Whitney U test. Correlation coefficient was evaluated by Spearman’s rank correlation test. Stab1, Stabilin-1; Stab2, Stabilin-2; T2DM, Type 2 diabetes mellitus; BMI, body mass index; LDL, low-density lipoprotein; HDL, high-density lipoprotein; SEM, standard error of mean.

In view of these findings, we further examined the association of Stab1 levels with classical CVD risk factors within this cohort. Spearman correlation analysis revealed a positive correlation between Stab1 and body mass index (BMI; r_s_ = 0.573, p < 0.0001; Fig. 1F). In contrast, Stab1 levels were negatively correlated with total cholesterol (r_s_ = − 0.342, p = 0.014; Fig. 1G), LDL cholesterol (r_s_ = − 0.331, p = 0.018; Fig. 1H) and HDL cholesterol (r_s_ = − 0.390, p = 0.005; Fig. 1I).

### Increased circulating levels of Stabilin-1 in T2DM patients with high atherosclerotic burden

Subsequently, we focused on the circulating levels of Stab1 & 2 and their ligands in T2DM patients (n=33), categorized based on the number of atherosclerotic plaques. As observed previously, TGFbI, Periostin and Stab2 were comparable between T2DM patients with low or high atherosclerotic plaque burden (low vs. high atherosclerotic burden: TGFbI: 543.6 ± 51.51 vs. 725.8 ±86.55 ng/ml, p = 0.13; Periostin: 103.6 ± 13.52 vs. 128.2 ±21.99 ng/ml, p = 0.655; Stab2: 3223 ± 309.7 vs. 3044 ± 263.1 pg/ml, p = 0.681; Fig. 1J, K, N), while Reelin levels exhibited a trend toward higher values in patients with high atherosclerotic burden (low vs. high atherosclerotic burden: Reelin: 1337 ± 210 vs. 2232 ± 431.4 pg/ml, p = 0.074; Fig. 1L). As also observed in the total cohort, Stab1 was also higher in T2DM patients with high atherosclerotic burden (low vs. high atherosclerotic burden: Stab1: 19.01 ± 1.815 vs. 29.11 ± 3.614 ng/ml, p = 0.045; Fig. 1M). Moreover, Stab1 retained the positive correlation with BMI (BMI; rs = 0.458, p = 0.009; Fig. 1O), as well as the negative correlations with total cholesterol (rs = − 0.432, p = 0.014; Fig. 1P) and LDL cholesterol (rs = − 0.473, p = 0.006; Fig. 1Q), when examining only T2DM patients. No correlation was detected between Stab1 and HDL (rs = − 0.216, p = 0.235; Fig. 1R).

## Discussion

The novel finding of this study is that Stabilin-1 may serve as a circulating marker of metabolic dysfunction and higher atherosclerotic burden in individuals with T2DM. The pathogenesis of atherosclerosis involves the interaction among endothelial dysfunction, lipid accumulation, oxidative stress and chronic inflammation^10,13,24^. Particularly, the infiltration of the arterial intima by LDL particles and their oxidative modification to form oxLDL lies at the core of atherogenesis, triggering endothelial injury and deformation and promoting local proinflammatory responses^7–9,11^. In the context of T2DM, hyperglycemia and dyslipidemia accelerate this cascade by promoting advanced glycation end products (AGEs) and impairing reverse cholesterol transport^17^. Stabilins, mainly Stab1, are crucial receptors that facilitate the endocytosis and the clearance of macromolecular waste products, including oxLDL^12,14,15,19^. In addition to its scavenging role, Stab1 also facilitates the removal of apoptotic cells, participates in leukocyte transmigration into inflammatory sites and has been found overexpressed in circulating monocytes in individuals with familial hypercholesterolemia^25^.

The role of Stab1 and Stab2 in atherogenesis has recently gained significant attention. Notably,, ApoE-Stab1-KO and ApoE-Stab2-KO mice were protected against both spontaneous and Western diet-induced atherogenesis^12^. Similarly, in ApoE-KO and Ldlr-KO mice, monoclonal antibodies against Stab1 or Stab2 markedly decreased Western diet-associated atherosclerosis^12^. Providing the clinical relevance of these results, we found that Stab1 was increased in patients with higher atherosclerotic plaque burden. In particular, we further found a positive correlation of Stab1 with BMI and negative correlations with total cholesterol and LDL cholesterol. While we did not detect an association between Stab2 and atherosclerotic burden, this could be probably attributed to the liver-specific expression of Stab2 in contrast to Stab1, which is strongly expressed in aortic endothelial cells and plaque macrophages, thereby probably affecting atherogenesis through additional local mechanisms^12^. Large prospective cohort studies are warranted to confirm the value of Stab1 as a potential prognostic factor for the development and progression of ASCVD.

Stabilins also exert their atheroprotective role through the removal of pro-inflammatory and pro-atherogenic ligands, including TGFbI, Periostin and Reelin^12^. TGFbI has been found to be expressed in smooth muscle cells and macrophages in human atherosclerotic plaques, while elimination of Periostin mitigates atherosclerosis in murine models through reducing macrophage recruitment^26,27^. Reelin loss also prevents atherosclerosis by decreasing leukocyte-endothelial cell adhesion and lesion macrophage accumulation^28^. In line with these findings, Reelin levels were elevated in diabetic patients with higher atherosclerotic burden in this study.

As mentioned, an important and novel finding of the present study is the inverse correlation of Stab1 levels with circulating cholesterol, LDL and HDL cholesterol. According to a preclinical study, there was no altered lipidomic profile between wild-type and Stab1 / Stab2-KO mice^12^. Similarly, bone marrow-specific knockout of Stab1 in hypercholesterolemic mice lipid did not affect blood levels or size and composition of atherosclerotic plaques^29^. Whether stabilins play a novel role as regulators of lipid metabolism in the context of diabetes remains to be examined in future mechanistic *in vivo* studies.

Limitations of this study include a relatively small sample size, which precludes further analyses, such as any potential effect by sex, as well as the use of multi-parametric regression models to examine the independent value of stabilins as markers of atherosclerotic burden. Another limitation is the cross-sectional study design, which does not allow us to examine the prognostic value of stabilins for atherosclerosis progression or major cardiovascular events.

In conclusion, this study indicates Stab1 as a promising circulating indicator of the atherosclerotic burden in T2DM patients. Stab1 complex and crucial role is highlighted by its involvement in clearing oxLDL and inflammatory mediators, combined with the positive correlation to BMI and negative correlation to total and LDL cholesterol. Comprehensive *in vivo* experiments in atherosclerosis models, together with large prospective cohort studies, are warranted to establish a causal link between stabilins and atherosclerosis development and progression, as well as to determine their prognostic value in ASCVD and other metabolic diseases.

## Competing interests

Authors declare that they have no competing interests

### Resource availability

#### Lead contact

Assoc. Prof. Antonios Chatzigeorgiou

#### Data and materials availability

The raw data of this study will be available upon reasonable request.

## References

1. Cho NH, Shaw JE, Karuranga S, Huang Y, da Rocha Fernandes JD, Ohlrogge AW, Malanda B. IDF Diabetes Atlas: Global estimates of diabetes prevalence for 2017 and projections for 2045. Diabetes Research and Clinical Practice. 2018;138:271–281.

2. Rawshani A, Rawshani A, Franzén S, Eliasson B, Svensson A-M, Miftaraj M, McGuire DK, Sattar N, Rosengren A, Gudbjörnsdottir S. Mortality and Cardiovascular Disease in Type 1 and Type 2 Diabetes. The New England Journal of Medicine. 2017;376(15):1407–1418.

3. Einarson TR, Acs A, Ludwig C, Panton UH. Prevalence of cardiovascular disease in type 2 diabetes: a systematic literature review of scientific evidence from across the world in 2007–2017. Cardiovascular Diabetology. 2018;17(1):83.

4. Martín-Timón I, Sevillano-Collantes C, Segura-Galindo A, del Cañizo-Gómez FJ. Type 2 diabetes and cardiovascular disease: Have all risk factors the same strength? World Journal of Diabetes. 2014;5(4):444– 470.

5. Ference BA, Ginsberg HN, Graham I, Ray KK, Packard CJ, Bruckert E, Hegele RA, Krauss RM, Raal FJ, Schunkert H, Watts GF, Boren J, Fazio S, Horton JD, Masana L, et al. Low-density lipoproteins cause atherosclerotic cardiovascular disease. 1. Evidence from genetic, epidemiologic, and clinical studies. A consensus statement from the European Atherosclerosis Society Consensus Panel. EUROPEAN HEART JOURNAL. 2017;38(32):2459–2472.

6. Low Wang CC, Hess CN, Hiatt WR, Goldfine AB. Clinical Update: Cardiovascular Disease in Diabetes Mellitus: Atherosclerotic Cardiovascular Disease and Heart Failure in Type 2 Diabetes Mellitus - Mechanisms, Management, and Clinical Considerations. Circulation. 2016;133(24):2459–2502.

7. Li J, Shangguan H, Chen X, Ye X, Zhong B, Chen P, Wang Y, Xin B, Bi Y, Zhu D. Advanced glycation end product levels were correlated with inflammation and carotid atherosclerosis in type 2 diabetes patients. Open Life Sciences. 2020;15(1):364–372.

8. Ménégaut L, Laubriet A, Crespy V, Leleu D, Pilot T, Van Dongen K, de Barros J-PP, Gautier T, Petit J-M, Thomas C, Nguyen M, Steinmetz E, Masson D. Inflammation and oxidative stress markers in type 2 diabetes patients with Advanced Carotid atherosclerosis. Cardiovascular Diabetology. 2023;22(1):248.

9. Bergheanu SC, Bodde MC, Jukema JW. Pathophysiology and treatment of atherosclerosis. Netherlands Heart Journal. 2017;25(4):231–242.

10. Goderis G, Vaes B, Mamouris P, Craeyveld E van, Mathieu C. Prevalence of Atherosclerotic Cardiovascular Disease, Heart Failure, and Chronic Kidney Disease in Patients with Type 2 Diabetes Mellitus: A Primary Care Research Network-based Study. Experimental and Clinical Endocrinology & Diabetes. 2021;130:447– 453.

11. Borén J, Chapman MJ, Krauss RM, Packard CJ, Bentzon JF, Binder CJ, Daemen MJ, Demer LL, Hegele RA, Nicholls SJ, Nordestgaard BG, Watts GF, Bruckert E, Fazio S, Ference BA, et al. Low-density lipoproteins cause atherosclerotic cardiovascular disease: pathophysiological, genetic, and therapeutic insights: a consensus statement from the European Atherosclerosis Society Consensus Panel. European Heart Journal. 2020;41(24):2313–2330.

12. Manta C-P, Leibing T, Friedrich M, Nolte H, Adrian M, Schledzewski K, Krzistetzko J, Kirkamm C, David Schmid C, Xi Y, Stojanovic A, Tonack S, de la Torre C, Hammad S, Offermanns S, et al. Targeting of Scavenger Receptors Stabilin-1 and Stabilin-2 Ameliorates Atherosclerosis by a Plasma Proteome Switch Mediating Monocyte/Macrophage Suppression. Circulation. 2022;146(23):1783–1799.

13. Rafieian-Kopaei M, Setorki M, Doudi M, Baradaran A, Nasri H. Atherosclerosis: Process, Indicators, Risk Factors and New Hopes. International Journal of Preventive Medicine. 2014;5(8):927–946.

14. Moore KJ, Freeman MW. Scavenger Receptors in Atherosclerosis. Arteriosclerosis, Thrombosis, and Vascular Biology. 2006;26(8):1702–1711.

15. van Berkel TJ, Out R, Hoekstra M, Kuiper J, Biessen E, van Eck M. Scavenger receptors: friend or foe in atherosclerosis? Current Opinion in Lipidology. 2005;16(5):525.

16. Rantakari P, Patten DA, Valtonen J, Karikoski M, Gerke H, Dawes H, Laurila J, Ohlmeier S, Elima K, Hübscher SG, Weston CJ, Jalkanen S, Adams DH, Salmi M, Shetty S. Stabilin-1 expression defines a subset of macrophages that mediate tissue homeostasis and prevent fibrosis in chronic liver injury. Proceedings of the National Academy of Sciences of the United States of America. 2016;113(33):9298– 9303.

17. Twarda-Clapa A, Olczak A, Białkowska AM, Koziołkiewicz M. Advanced Glycation End-Products (AGEs): Formation, Chemistry, Classification, Receptors, and Diseases Related to AGEs. Cells. 2022;11(8):1312.

18. Leibing T, Riedel A, Xi Y, Adrian M, Krzistetzko J, Kirkamm C, Dormann C, Schledzewski K, Goerdt S, Géraud C. Deficiency for scavenger receptors Stabilin-1 and Stabilin-2 leads to age-dependent renal and hepatic depositions of fasciclin domain proteins TGFBI and Periostin in mice. Aging Cell. 2023;22(9):e13914.

19. Li R, Oteiza A, Sørensen KK, McCourt P, Olsen R, Smedsrød B, Svistounov D. Role of liver sinusoidal endothelial cells and stabilins in elimination of oxidized low-density lipoproteins. American Journal of Physiology - Gastrointestinal and Liver Physiology. 2011;300(1):G71–G81.

20. Pandey E, Nour AS, Harris EN. Prominent Receptors of Liver Sinusoidal Endothelial Cells in Liver Homeostasis and Disease. Frontiers in Physiology. 2020;11.

21. Levey AS, Stevens LA, Schmid CH, Zhang YL, Castro AF, Feldman HI, Kusek JW, Eggers P, Van Lente F, Greene T, Coresh J, CKD-EPI (Chronic Kidney Disease Epidemiology Collaboration). A new equation to estimate glomerular filtration rate. Annals of Internal Medicine. 2009;150(9):604–612.

22. Chatzigeorgiou A, Mitroulis I, Chrysanthopoulou A, Legaki A-I, Ritis K, Tentolouris N, Protogerou AD, Koutsilieris M, Sfikakis PP. Increased Neutrophil Extracellular Traps Related to Smoking Intensity and Subclinical Atherosclerosis in Patients with Type 2 Diabetes. Thrombosis and Haemostasis. 2020;120:1587–1589.

23. Vlachogiannis NI, Legaki A-I, Kassi E, Mikelis CM, Tentolouris N, Sfikakis PP, Protogerou AD, Chatzigeorgiou A. Association of Circulating Robo4 with Obesity, Hypertension and Atherosclerotic Plaque Burden. Thrombosis and Haemostasis. 2024.

24. Ren H, Zhao,Lijun, Zou,Yutong, Wang,Yiting, Zhang,Junlin, Wu,Yucheng, Zhang,Rui, Wang,Tingli, Wang,Jiali, Zhu,Yitao, Guo,Ruikun, Xu,Huan, Li,Lin, Cooper,Mark E., and Liu F. Association between atherosclerotic cardiovascular diseases risk and renal outcome in patients with type 2 diabetes mellitus. Renal Failure. 2021;43(1):477–487.

25. Brochériou I, Maouche S, Durand H, Braunersreuther V, Naour GL, Gratchev A, Koskas F, Mach F, Kzhyshkowska J, Ninio E. Antagonistic regulation of macrophage phenotype by M-CSF and GM-CSF: Implication in atherosclerosis. Atherosclerosis. 2011;214(2):316–324.

26. O’Brien ER, Bennett KL, Garvin MR, Zderic TW, Hinohara T, Simpson JB, Kimura T, Nobuyoshi M, Mizgala H, Purchio A, Schwartz SM. Beta ig-h3, a transforming growth factor-beta-inducible gene, is overexpressed in atherosclerotic and restenotic human vascular lesions. Arteriosclerosis, Thrombosis, and Vascular Biology. 1996;16(4):576–584.

27. Schwanekamp JA, Lorts A, Vagnozzi RJ, Vanhoutte D, Molkentin JD. Deletion of Periostin Protects Against Atherosclerosis in Mice by Altering Inflammation and Extracellular Matrix Remodeling. Arteriosclerosis, Thrombosis, and Vascular Biology. 2016;36(1):60–68.

28. Ding Y, Huang L, Xian X, Yuhanna IS, Wasser CR, Frotscher M, Mineo C, Shaul PW, Herz J. Loss of Reelin protects against atherosclerosis by reducing leukocyte-endothelial adhesion and lesion macrophage accumulation. Science signaling. 2016;9(419):ra29.

29. Nahon JE, Hoekstra M, Hulst S van, Manta C, Goerdt S, Geerling JJ, Géraud C, Eck MV. Hematopoietic Stabilin-1 deficiency does not influence atherosclerosis susceptibility in LDL receptor knockout mice. Atherosclerosis. 2019;281:47–55.

